# An expanded palette of ATP sensors for subcellular and multiplexed imaging

**DOI:** 10.1101/2023.05.01.538722

**Authors:** Ayse Z. Sahan, Eventine Youngblood, Siddharth Das, Danielle L. Schmitt, Jin Zhang

## Abstract

Genetically encoded fluorescent biosensors that detect changes in ATP levels in live cells have enabled the discovery of novel roles for ATP in cellular processes and signaling. Many of these available ATP biosensors have a limited dynamic range, or have ATP affinities that are not suitable for sensing the physiological concentrations of ATP in mammalian cells. To address these limitations, we developed a FRET-based ATP biosensor with enhanced dynamic range and signal to noise ratio, eATeam. Using eATeam, we uncovered distinct spatiotemporal dynamics of ATP changes upon inhibition of cellular energy production. We also developed dimerization-dependent GFP and RFP-based ATP biosensors with enhanced dynamic ranges compared to the current standard in the field. Using the single-color ATP biosensors, we visualized the complex interplay between AMPK activity, ATP, lactate, and calcium by multiplexed imaging in single cells. This palette of ATP sensors expands the toolbox for interrogating subcellular ATP regulation and metabolic signaling in living cells.

## Introduction

Adenosine 5’-triphosphate (ATP) is an energy storage molecule of the cell that is crucial to nearly all cellular processes and functions, including muscle contraction^1^, neurotransmission,^2^ cell motility^3^, and purinergic signaling^4^. As a central metabolite of the cell, ATP is highly involved in metabolic signaling pathways that mediate cellular function in response to dynamic changes in nutrient levels. This enables cells to finely tune signaling and gene expression to the metabolic state of the cell^5^. For instance, AMP-activated protein kinase (AMPK) is a metabolite sensor that responds to changing glucose and energy status. Under low ATP conditions, AMPK is rapidly activated to redirect metabolism towards catabolic processes and release stored energy^6^. Spatial regulation of AMPK is critical for achieving high signaling specificity^7^. Previous studies have discussed the presence of ATP at mitochondria, the endoplasmic reticulum, and the nucleus^8–10^, but a more comprehensive understanding of subcellular pools of ATP is required to understand the dynamic interplay between ATP and spatial regulation of signaling pathways such as AMPK. Given its importance to a variety of cellular processes, understanding the spatiotemporal organization and regulation of ATP can provide critical insights to how this energy currency coordinates and balances metabolic and signaling processes with high specificity.

Significant advances have been made in developing fluorescent protein-based biosensors for detecting changes in ATP levels in live cells. The first generation ATP biosensor, ATeam1.03 (Adenosine 5-Triphosphate indicator based on Epsilon subunit for Analytical Measurements), is a Fluorescence Resonance Energy Transfer (FRET)-based biosensor that directly binds ATP^11^. ATeam1.03 incorporates the epsilon subunit of the F_0_F_1_-ATP synthase from *Bacillus subtilis* as the ATP sensing subunit of the biosensor. On the N- and C-termini of this sensing subunit are mseCFP (monomeric super enhanced cyan fluorescent protein) and circularly permutated (cp) mVenus, respectively. The sensing subunit has two alpha helices on the C-terminus which directly bind ATP and interact with an N-terminal β-sandwich, leading to a conformational change that is favorable for FRET which can be detected as an increase in the yellow/cyan emission ratio. Since the introduction of ATeam1.03, several sensors incorporating the same or similar sensing subunits have been developed with various K_d_ values for use in yeast, bacteria and mammalian cells^12–15^. Fluorescent protein-based ATP biosensors like ATeam1.03, the MaLion series^12^, Queen^15^, and iATPSnFR^13^ have elucidated changes in ATP levels by live-cell imaging. The current standard in the field is still ATeam1.03, which exhibits an optimal dissociation value (K_d_ = 3.3 mM) for detecting changes in intracellular ATP concentrations, which range from 0.5 to 5 mM^16^, and has been validated for use in mammalian cells. ATeam1.03 has, to this point, been crucial for demonstrating the utility of fluorescent protein-based biosensors in characterizing the roles of ATP. However, there are certain drawbacks which we aimed to address in this study. The dynamic range of ATeam1.03 is limited (∼15-20% response to 2-DG in mammalian cells reported)^11^, and currently available ATP biosensors for use in mammalian cells perform similarly. A stand-out in the field is the excitation ratiometric yellow sensor PercevalHR, which reports on the ATP:ADP ratio in live cells and exhibits a reported 70% response to 2-DG treatment^17^. However, there is still a need for an ATP biosensor with improved dynamic range and signal to noise ratio, as the capabilities of current sensors may limit their utility for probing minute changes in subcellular pools of ATP. In addition to enhancing dynamic range, we set out to develop an expanded palette of single-color ATP biosensors based on the design of ATeam1.03 to augment multiplexing capabilities. Expanding the current toolbox by developing ATP biosensors with novel reporting units and red-shifting fluorescence will greatly enhance multiplexing capabilities, freeing up spectral space for co-imaging with other biosensors.

Here, we report a palette of improved FRET-based and single-color ATP biosensors that enable visualization of distinct subcellular ATP dynamics and multiplexed imaging of ATP. We incorporate a design that is new to ATP biosensors, dimerization dependent fluorescent proteins, for achieving single-color imaging of changes in cellular ATP levels. We also present a red ATP biosensor that is further red-shifted compared to the available red ATP biosensor MaLionR^12^.

## Results

### Development of an improved Y/C FRET-based ATP biosensor

ATeam1.03 incorporates the epsilon subunit of the *B. subtilis* F_0_F_1_-ATP synthase sandwiched between a cyan (mseCFP) and a yellow (cp mVenus[E172]) fluorescent protein pair (**Figure 1A**). Upon binding ATP, the sensing subunit undergoes a conformational change that leads to an increase in the Y/C emission ratio. We aimed to achieve improved dynamic range of the ATeam1.03 sensor without changing the sensing subunit, which has been well-characterized for its affinity, specificity, and sensitivity for detecting cellular ATP levels^11^. Our first strategy was to switch out the FRET pair in ATeam1.03 (mseCFP and cpmVenus [E172]) with alternative yellow and cyan fluorescent protein pairs. To this end, we screened 3 fluorescent protein pairs that are commonly used in FRET-based biosensors:^18–20^ mTurquoise2 and cpmVenus E172, mCerulean and cpVenus E172, and mCerulean and YPet. HEK293T cells transiently expressing each construct were imaged for 25 minutes following treatment with 40 mM of 2-deoxyglucose (2-DG), an inhibitor of glycolysis that rapidly decreases intracellular ATP concentrations. Upon 2-DG treatment (n = 50), ATeam1.03 emission ratio decreased by 8.5 ± 0.9% on average compared to the vehicle control (DMSO, n = 34) (**Figure 1B, C**). We identified an improved FRET-based ATP biosensor, enhanced ATeam (eATeam), which utilizes mCerulean and YPet as the FRET pair (**Figure 1D**). 2-DG treatment (n = 27) induced an 12.5 ± 0.9% change (p-value < 0.0001) in the eATeam emission ratio, whereas the vehicle control had no effect (n = 26) (**Figure 1E, F**). eATeam exhibited a 40.1 ± 10.3% (n = 26, p-value < 0.001) improved response compared to ATeam1.03 (n = 34) in HEK293T cells (**Figure 1G**). In addition to an improved response, eATeam exhibited a faster time to half of the maximal response (t_1/2_) than ATeam1.03 (**Figure 1H**, ATeam1.03, 11.1 ± 0.5 min, eATeam, 7.1 ± 0.6 min, p-value < 0.0001). This faster response time may indicate improved kinetics of eATeam and may be a consideration in experimental design for probing ATP. The signal to noise ratio (SNR) of eATeam (44.0 ± 3.3) was also significantly improved compared to ATeam1.03 (29.2 ± 3.7) (**Figure 1I**, p-value < 0.01). The ATP biosensor variants were also tested in MEF cells (**Supplementary Figure 1**). eATeam exhibited a 103 ± 17.5% (p-value < 0.0001) higher response to 2-DG treatment (n = 10) in MEF cells compared to ATeam1.03 (n = 19) (**Supplementary Figure 1E**), faster t_1/2_ (**Supplementary Figure 1F**, ATeam1.03, 10.6 ± 0.4 min, eATeam, 5.0 ± 0.7 min, p-value < 0.0001), and higher signal to noise ratio (**Supplementary Figure 1G**, ATeam1.03, 41.8 ± 5.3, eATeam, 71.3 ± 11.9, p-value < 0.05), indicating that the observed improvement in response is not cell type specific. Ultimately, we found that eATeam exhibits a greater response to 2-DG and a higher signal to noise ratio compared to ATeam1.03.

**Figure 1.**
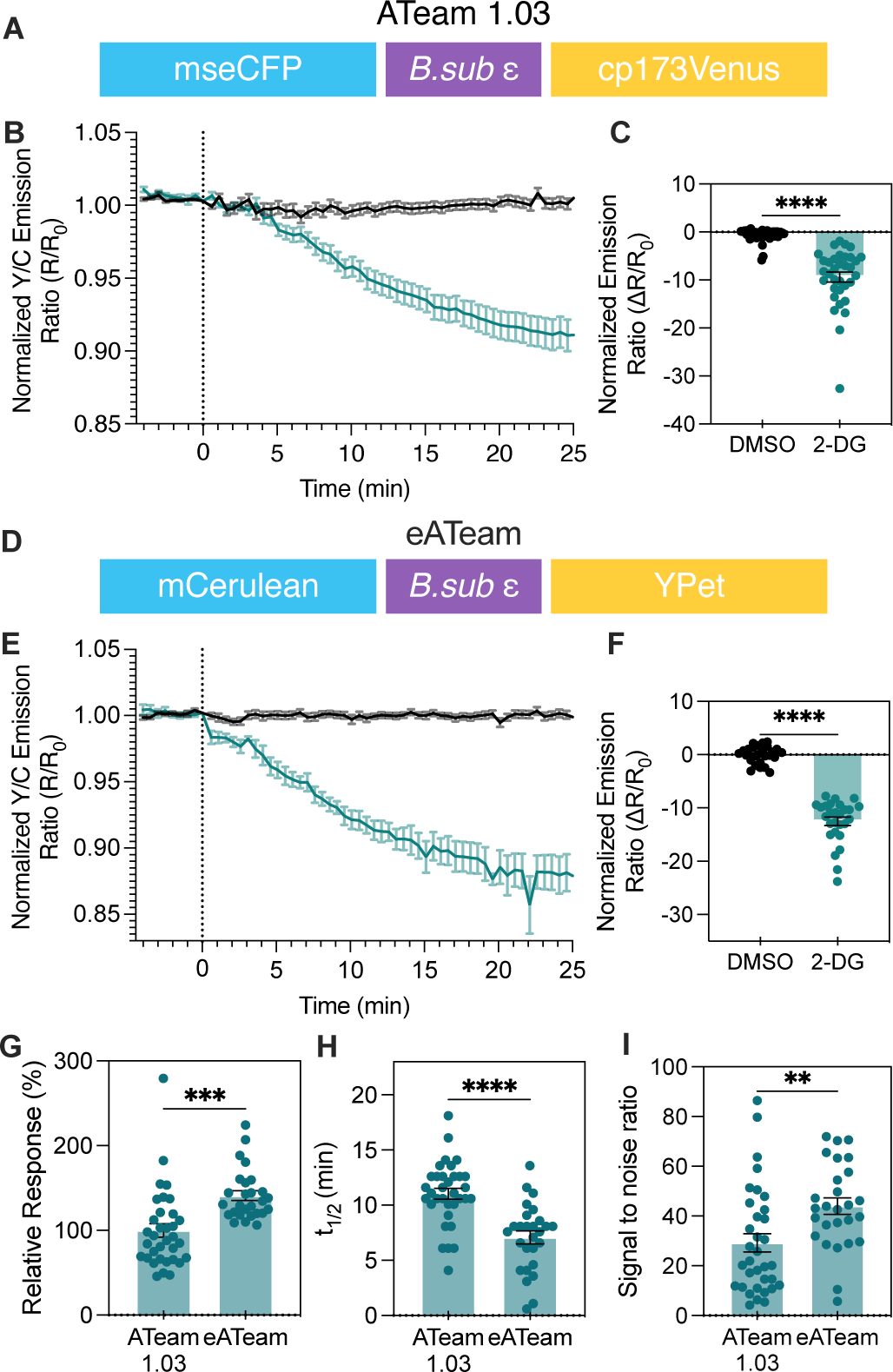
Development of an improved Y/C FRET-based ATP biosensor derived from ATeam1.03. **(A)** Domain structure of ATeam1.03. **(B)** Averaged time course traces of normalized emission ratio (yellow/cyan) of ATeam1.03 in HEK293T cells treated with DMSO (black, vehicle control) or 2-DG (blue). Data were from at least three independent experiments. Shaded area indicates standard error of the mean (SEM). **(C)** Bar graph shows the average response at 25 minutes after DMSO (n = 50) or 2-DG (n = 34) addition. Unpaired student’s t-test (two-tailed) is used. ****, p<0.0001. **(D)** Domain structure of eATeam. **(E)** Averaged time course traces of normalized emission ratio (yellow/cyan) of eATeam in HEK293T cells treated with DMSO (black, vehicle control) or 2-DG (blue) treatment in HEK293T cells. Data were from at least three independent experiments. Shaded area indicates SEM. **(F)** Bar graph shows the average response at 25 minutes after DMSO (n = 26) or 2-DG (n = 27) addition. Unpaired student’s t-test (two-tailed) is used. ****, p < 0.0001. **(G)** Comparison of 2-DG induced responses of ATeam1.03 (n = 34) and eATeam (n=27). Calculated by dividing each response value by the mean response of ATeam1.03. Bar graph shows the average response at 25 minutes after 2-DG addition and error bars represent SEM. Unpaired student’s t-test (two-tailed) is used. ***, p < 0.001. **(H)** Direct comparison of ATeam1.03 (n = 34) and eATeam (n=27) time to half minimum response (t_1/2_) in minutes. Bar graph shows the average t_1/2_ and error bars represent SEM. Unpaired student’s t-test (two-tailed) is used. ****, p < 0.0001. **(I)** Direct comparison of ATeam1.03 (n = 34) and eATeam (n=27) signal to noise ratio. Bar graph shows the average ratio and error bars represent SEM. Unpaired student’s t-test (two-tailed) is used. **, p < 0.01.

### Characterization of subcellular ATP levels using the improved FRET-based ATP biosensor eATeam

After identifying an improved FRET-based ATP biosensor (eATeam), we aimed to demonstrate its use in characterizing subcellular pools of ATP. Cellular energy is largely produced from glucose or metabolites fed into glucose metabolism, which is used in three distinct and subsequent processes to produce ATP: glycolysis, tricaboxylic acid cycle (TCA), and oxidative phosphorylation. During glycolysis, glucose is catabolized to produce pyruvate and a relatively small amount of ATP. Acetyl coenzyme A is generated from pyruvate and used to produce NADH and FADH_2_ through TCA. Finally, NADH and FADH_2_ are used in the electron transport chain to maintain a proton gradient across the inner mitochondrial membrane so that mitochondrial ATP synthase can produce large amounts of ATP. All of this occurs around or inside of mitochondria, which then supplies ATP to other locations in the cell^21^. By using CCCP to uncouple the electron transport chain and inhibit oxidative phosphorylation and 2-DG to inhibit glycolysis, we aimed to distinguish ATP depletion characteristics at various locations across the cell.

Like ATeam1.03, untargeted eATeam is excluded from the nucleus (**Figure 2A, B**), and was therefore used to probe cytosolic ATP. eATeam was also targeted to the plasma membrane (**Figure 2F, G**), the outer mitochondrial membrane (**Figure 2K, L**), and the nucleus (**Figure 2P, Q**) by the addition of Lyn^22^, dAKAP^23^, and H2A^24^ subcellular targeting motifs, respectively, to the N-terminus of eATeam (sequences listed in **Methods and Materials** section). HEK293T cells expressing each subcellularly localized biosensor were imaged upon addition of DMSO (vehicle control), 5 nM CCCP (mitochondrial uncoupler) to inhibit oxidative phosphorylation, or 40 mM 2-DG to inhibit glycolysis.

**Figure 2.**
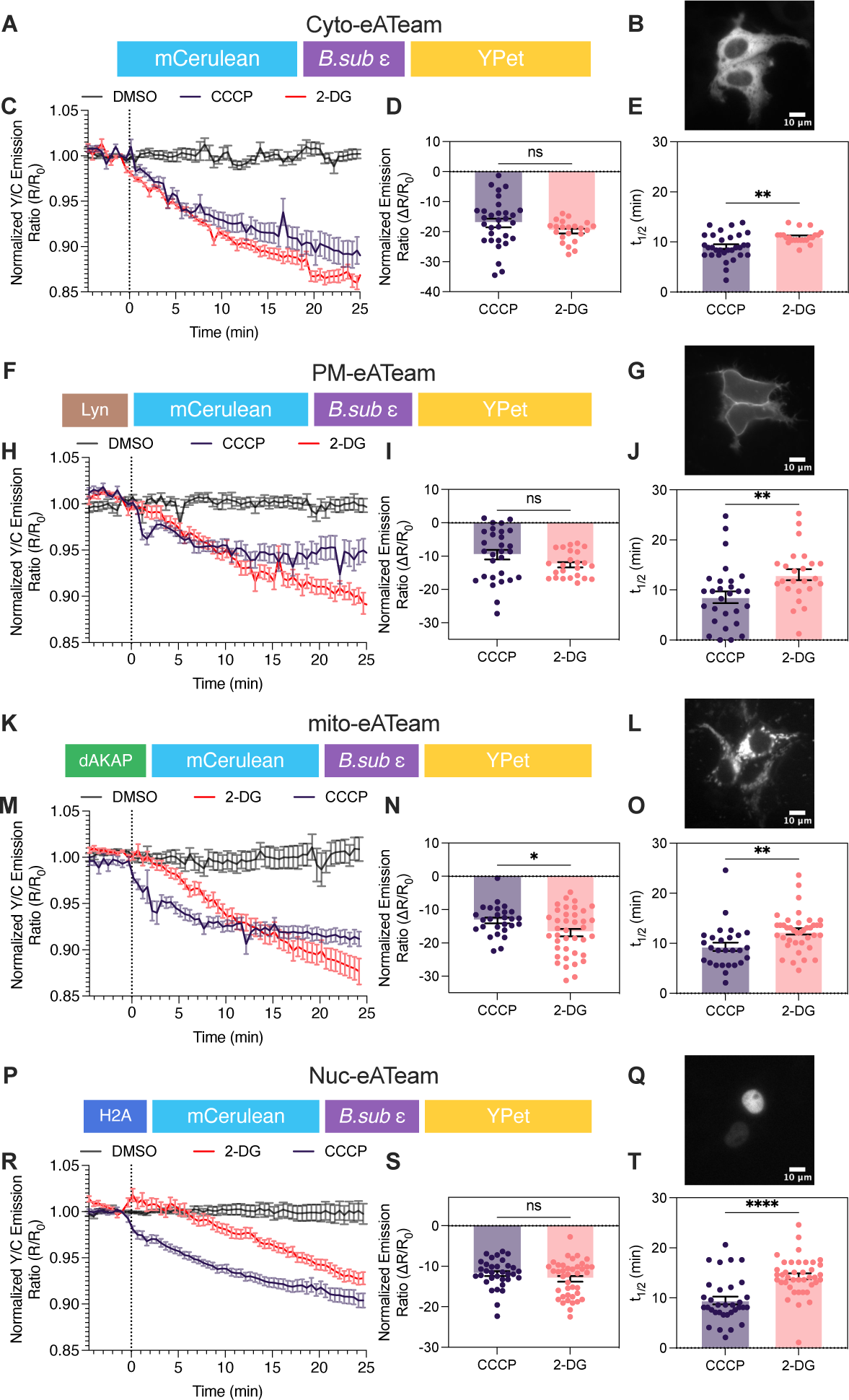
Characterization of subcellular ATP levels using the improved FRET-based ATP biosensor eATeam. **(A)** Domain structure of cytosolic eATeam. **(B)** Representative fluorescence images of HEK293T cells expressing cytosolic eATeam. Scale bar = 10 µm. **(C)** Averaged time course traces of normalized emission ratio (yellow/cyan) of cyto-eATeam in HEK293T cells treated with DMSO (black, vehicle control), 2-DG (pink), or CCCP (purple) treatment. Data were from at least three independent experiments. Shaded area indicates standard error of the mean (SEM). **(D)** Summary of 2-DG (n = 22) or CCCP (n =31) induced cyto-eATeam responses in (C). Bar graph shows the average response at 25 minutes after drug addition and error bars represent SEM. Unpaired student’s t-test (two-tailed) is used. ns, p is not significant. **(E)** Summary of t_1/2_ of cyto-eATeam responses induced by 2-DG (n = 22), or CCCP (n =31). Bar graph shows the average t_1/2_ and error bars represent SEM. Unpaired student’s t-test (two-tailed) is used. **, p < 0.01. **(F)** Domain structure of plasma membrane targeted PM-eATeam. **(G)** Representative fluorescence images of HEK293T cells expressing PM-eATeam. Scale bar = 10 µm. **(H)** Averaged time course traces of normalized emission ratio (yellow/cyan) of PM-eATeam in HEK293T cells treated withDMSO (black, vehicle control), 2-DG (pink), or CCCP (purple) treatment. Data were from at least three independent experiments. Shaded area indicates standard error of the mean (SEM). **(I)** Summary of 2-DG (n = 24), or CCCP (n =29) induced PM-eATeam responses in (H). Bar graph shows the average response 25 minutes after drug addition and error bars represent SEM. Unpaired student’s t-test (two-tailed) is used. ns, p is not significant. **(J)** Summary of t_1/2_ of PM-eATeam responses induced by 2-DG (n = 24), or CCCP (n =29). Bar graph shows the average t_1/2_ and error bars represent SEM. Unpaired student’s t-test (two-tailed) is used. **, p < 0.01. **(K)** Domain structure of outer mitochondrial membrane targeted mito-eATeam. **(L)** Representative fluorescence images of HEK293T cells expressing mito-eATeam. Scale bar = 10 µm. **(M)** Averaged time course traces of normalized emission ratio (yellow/cyan) of mito-eATeam in HEK293T cells treated with DMSO (black, vehicle control), 2-DG (pink), or CCCP (purple) treatment. Data were from at least three independent experiments. Shaded area indicates standard error of the mean (SEM). **(N)** Summary of 2-DG (n = 24), or CCCP (n =29) induced mito-eATeam responses in (M). Bar graph shows the average response 25 minutes after drug addition and error bars represent SEM. Unpaired student’s t-test (two-tailed) is used. *, p < 0.05. **(O)** Summary of t_1/2_ of mito-eATeam responses induced by 2-DG (n = 24), or CCCP (n =29). Bar graph shows the average t_1/2_ and error bars represent SEM. Unpaired student’s t-test (two-tailed) is used. **, p < 0.01. **(P)** Domain structure of nucleus targeted Nuc-eATeam. **(Q)** Representative fluorescence images of HEK293T cells expressing Nuc-eATeam. Scale bar = 10 µm. **(R)** Averaged time course traces of normalized emission ratio (yellow/cyan) of Nuc-eATeam in HEK293T cells treated with DMSO (black, vehicle control), 2-DG (pink), or CCCP (purple) treatment. Data were from at least three independent experiments. Shaded area indicates standard error of the mean (SEM). **(S)** Summary of 2-DG (n = 24), or CCCP (n =29) induced Nuc-eATeam responses in (R). Bar graph shows the average response 25 minutes after drug addition and error bars represent SEM. Unpaired student’s t-test (two-tailed) is used. ns, p is not significant. **(T)** Summary of t_1/2_ of H2A-eATeam responses induced by 2-DG (n = 24), or CCCP (n =29). Bar graph shows the average t_1/2_ and error bars represent SEM. Unpaired student’s t-test (two-tailed) is used. ****, p<0.0001.

In the cytosol, the eATeam (cyto-eATeam) response induced by 2-DG (n = 22) or CCCP (n = 31) was not significantly different (**Figure 2C, D**, 2-DG, −19.8 ± 0.8 %, CCCP, −17.1 ± 1.4 %, p-value > 0.05). However, the t_1/2_ of cyto-eATeam response to 2-DG was significantly longer than with CCCP treatment (**Figure 2E**, CCCP, 9.1 ± 0.5 mins, 2-DG, 11.0 ± 0.3 min, p-value < 0.01), indicating a slower cytosolic ATP depletion upon inhibition of glycolysis. This trend was consistent with other pools of ATP (**Figure 2J, O, T**). Although glycolysis generates ATP more rapidly than oxidative phosphorylation, oxidative phosphorylation has a larger impact on the total cellular ATP^25^. This may account for the rapid depletion in ATP levels upon CCCP treatment. Interestingly, we observed distinct eATeam response amplitudes to 2-DG and CCCP at the mitochondria but not at the cytosol (**Figure 2D**), plasma membrane (**Figure 2I**), and nucleus (**Figure 2S**). Mitochondrial eATeam exhibited a larger response to 2-DG (n = 39) than CCCP (n = 27) (**Figure 2N**, CCCP, −13.3 ± 0.9, 2-DG, −16.9 ± 1.1, p-value < 0.05). Such subcellular differences in ATP depletion may reflect distinct coupling mechanisms between energy production and delivery of ATP to various locations across the cell. These observations also raise questions about differing energy dependencies across organelles, which is an interesting avenue for future exploration using subcellular eATeam.

We were specifically interested in probing changes in nuclear ATP, since it is a distinct and separate pool of ATP that is relatively understudied. It has been reported that ATP is supplied to the nucleus through mitochondrial oxidative phosphorylation^26^. It comes to reason that inhibiting mitochondrial energy production will lead to nuclear ATP depletion. Indeed, 2-DG and CCCP treatment significantly reduced Nuc-eATeam response (**Figure 2R, S**). Interestingly, we found that the nuclear eATeam response to 2-DG was significantly slower than the cytosolic response (**Supplementary Figure 2B**, nuclear t_1/2,_ 14.3 ± 0.6 min, cytosolic t_1/2_, 11.0 ± 0.3 min, p-value < 0.05), whereas the CCCP response was comparable to all other subcellular compartments. This hints at distinct mechanisms of ATP delivery to the nucleus originating from glycolysis or oxidative phosphorylation. Taken together, eATeam exhibits a dynamic range and sensitivity sufficient to characterize subcellular ATP levels and underscores the distinct spatiotemporal regulation of ATP.

### Developing an expanded palette of single-color dimerization-dependent fluorescent proteins (ddFP)-based ATP biosensors

We next sought to develop single-color ATP biosensors with improved dynamic ranges to achieve simultaneous imaging of subcellular pools of ATP. Our strategy was to use dimerization dependent fluorescent proteins (ddFPs), which are minimally fluorescent in the non-dimerized state, and exhibit increased fluorescence upon dimerization^27, 28^. We developed ddRFP (**Figure 3A**) and ddGFP (**Figure 3D**)-based intensiometric ATP biosensors utilizing the same sensing subunit and linker regions as ATeam1.03 and eATeam. The red biosensor, R-ATeam, exhibited an average decrease in response to 2-DG of 15.0 ± 2.1% (n = 28, p-value < 0.0001) (**Figure 3B, C**), which was comparable to eATeam and greater than ATeam1.03 (**Supplementary Figure 3C**). We also observed that the R-ATeam response to 2-DG is comparable to another red ATP biosensor, MaLionRed^12^, in terms of both amplitude and kinetics (**Supplementary Figure 3C, D**, p-value > 0.05).

**Figure 3.**
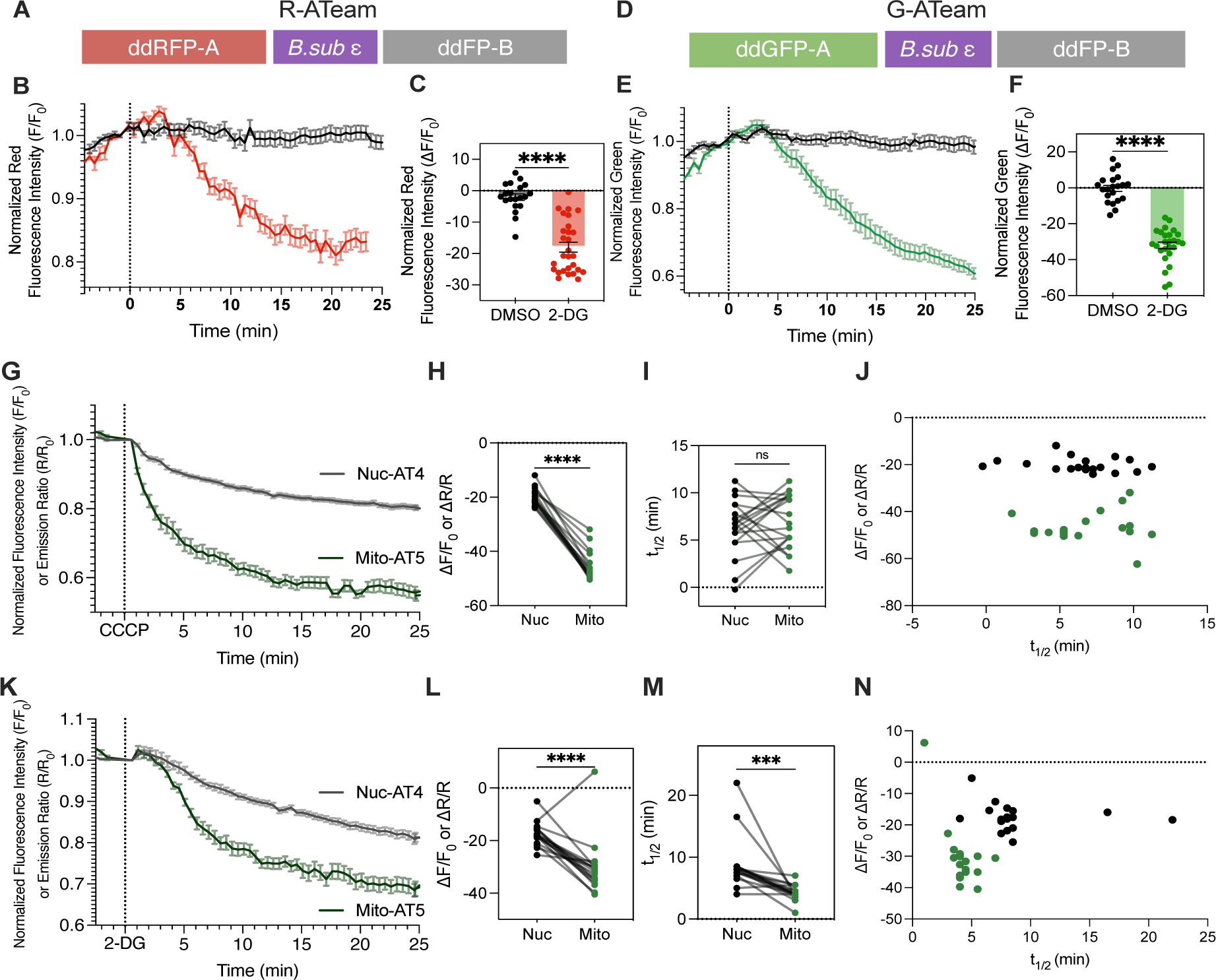
Development of an expanded palette of single-color ddFP-based ATP sensors. **(A)** Domain structure of R-ATeam, a red ATP sensor. **(B)** Averaged time course traces of normalized red fluorescence intensity of R-ATeam induced by DMSO (black, vehicle control) or 2-DG (red) treatment. Data were from at least three independent experiments. Shaded area indicates standard error of the mean (SEM). **(C)** Summary of DMSO (n = 20) or 2-DG (n = 28) induced R-ATeam responses in (B). Bar graph shows the average response at 25 minutes after treatment. Error bar represents SEM. Unpaired student’s t-test (two-tailed) is used. ****, p < 0.0001. **(D)** Domain structure of G-ATeam, a green ATP sensor. **(E)** Averaged time course traces of normalized green fluorescence intensity of G-ATeam induced by DMSO (black, vehicle control) or 2-DG (green) treatment. Data were from at least three independent experiments. Shaded area indicates standard error of the mean (SEM). **(F)** Summary of DMSO (n = 21) or 2-DG (n = 26) induced G-ATeam responses in (E). Bar graph shows the average response at 25 minutes after treatment. Error bar represents SEM. Unpaired student’s t-test (two-tailed) is used. ****, p < 0.0001. **(J)** Averaged time course traces of normalized emission ratio (yellow/cyan) of Nuc-eATeam (black) or normalized red fluorescence intensity of mito-R-ATeam (green) after CCCP treatment in HeLa cells co-expressing both sensors. Data were from at least three independent experiments. Shaded area indicates standard error of the mean (SEM). **(H)** Response of Nuc-eATeam or Mito-R-ATeam (n = 19) to CCCP treatment. Lines are connecting responses observed in the same cell. Paired student’s t-test (two-tailed) is used. ****, p < 0.0001. **(I)** t_1/2_ of Nuc-eATeam or Mito-R-ATeam (n = 19) response after CCCP treatment. Lines are connecting T one-halves obtained from the same cell. Paired student’s t-test (two-tailed) is used. ns, p is not significant. **(J)** Response of Nuc-eATeam (black) and Mito-R-ATeam to CCCP plotted against corresponding t_1/2_. **(K)** Averaged time course traces of normalized emission ratio (yellow/cyan) of Nuc-eATeam (black) or normalized red fluorescence intensity of mito-R-ATeam (green) after 2-DG treatment in HeLa cells co-expressing both sensors. Data were from at least three independent experiments. Shaded area indicates standard error of the mean (SEM). **(L)** Response of Nuc-eATeam or Mito-R-ATeam (n = 19) to 2-DG treatment. Lines are connecting responses observed in the same cell. Paired student’s t-test (two-tailed) is used. ****, p < 0.0001. **(M)** t_1/2_ of Nuc-eATeam or Mito-R-ATeam (n = 19) response after CCCP treatment. Lines are connecting T one-halves obtained from the same cell. Paired student’s t-test (two-tailed) is used. ***, p < 0.001. **(N)** Response of Nuc-eATeam (black) and Mito-R-ATeam to 2-DG plotted against corresponding t_1/2_.

We attempted to improve the dynamic range of the single-color ATP biosensors using R-ATeam as the template. We first incorporated successively longer flexible linkers on either end of the ATP sensing subunit to increase the distance between the two FPs (RFP-A and FP-B) in the ATP unbound state and maximize the change in sensor response (**Supplementary Figure 3A**, R-ATeam.2-.6). We also altered the amino acid sequence of the ATP sensing subunit, which has been extensively characterized previously^29–32^. Previously, Hires et al has shown that serial deletions of amino acids from either end of the sensing subunit of FRET biosensors have resulted in improved dynamic range^33^. We designed R-ATeam variants with 5-15 amino acid deletion from the N-terminus of the ATP sensing subunit (**Supplementary Figure 3A**, R-ATeam.7-.10). Our third and final strategy was to incorporate tandem copies of the FP-B fragment onto the C-terminus of the construct (**Supplementary Figure 3A**, R-ATeam 2x FP-B). Ultimately, we did not observe significantly improved response to 2-DG in any of the tested R-ATeam variants (**Supplementary Figure 3B**, p-values > 0.05).

To expand the toolbox of single-color ddFP-based ATP biosensors, we also developed a variant incorporating ddGFP that was designed with the same strategy as R-ATeam. The green biosensor, G-ATeam, was expressed in HEK293T cells and imaged upon treatment with 2-DG. G-ATeam had a further enhanced response to 2-DG treatment (n = 26) compared with ATeam1.03, with a 32.1 ± 1.9 % decrease in normalized GFP emission (**Figure 3E, F**, p-value < 0.0001). The relative response of G-ATeam to 2-DG treatment was, on average, 150.4 ± 14.3 % higher than ATeam1.03. G-ATeam is the best performing ATP biosensor we present (**Supplementary Figure 3C**, p-value < 0.0001).

We next demonstrated the use of these single-color ATP biosensors in co-imaging applications. R-ATeam was targeted to the outer mitochondrial membrane by a dAKAP targeting sequence fused to its N-terminus (Mito-R-ATeam). To simultaneously identify changes in nuclear and mitochondrial ATP upon CCCP or 2-DG treatment, we imaged HeLa cells co-expressing Nuc-eATeam (nucleus targeted FRET-based ATP biosensor) and Mito-R-ATeam (outer mitochondrial membrane targeted red ATP biosensor). CCCP treatment (n = 19) induced a reduction in both Mito-R-ATeam and Nuc-eATeam signals (**Figure 3G, H**). t_1/2_ of Nuc-eATeam response and Mito-R-ATeam response to CCCP were comparable (p-value > 0.05). 2-DG treatment, on the other hand, induced distinct kinetics for nuclear and mitochondrial ATP biosensors. Mito-R-ATeam had, in general, a faster response to 2-DG than Nuc-eATeam within the same cell (analyzed by paired t-test) (**Figure 3M**, Mito-R-ATeam, 4.2 ± 0.3, Nuc-eATeam, 8.8 ± 1.0, p-value < 0.001). We observed this trend in cells expressing each biosensor separately as well (**Supplementary Figure 2B**), although co-imaging allowed better comparison of the kinetics of ATP changes in two different locations despite cell-to-cell heterogeneity. Furthermore, there was greater variation in the t_1/2_ of Nuc-eATeam upon 2-DG treatment than Mito-R-ATeam (**Figure 3N**). We observed distinct kinetics of nuclear and mitochondrial ATP depletion upon glycolytic or oxidative phosphorylation inhibition. Using the single-color Mito-R-ATeam with the FRET-based Nuc-eATeam, we were able to probe changes in the nuclear and mitochondrial pools of ATP simultaneously within single cells and observe cell-to-cell heterogeneity in response.

### Multiplexed imaging of ATP with AMPK activity and other cellular analytes

Lastly, we demonstrated the utility of the single-color ATP biosensors in multiplexed imaging experiments with other biosensors. AMPK is a major stress and energy sensor of the cell and is closely tuned to fluctuations in ATP^34^. Importantly, mitochondria represent a key signaling node for AMPK^6^. Previously, ExRai-AMPKAR, a ratiometric green AMPK activity biosensor, was targeted to the outer mitochondrial membrane to probe dynamics of mitochondrial AMPK activity^7^. We therefore co-expressed mitochondria-targeted ExRai-AMPKAR (mito-ExRai-AMPKAR) and mito-R-ATeam to simultaneously probe the effect of 2-DG or CCCP on ATP levels and AMPK activity at the mitochondria of HeLa cells (**Figure 4A**). 2-DG (n = 20) and CCCP (n = 24) both increased the mito-ExRai-AMPKAR excitation ratio (480/405). This was consistently accompanied by a decrease in mitochondrial ATP (reduced mito-R-ATeam RFP intensity) (**Figure 4B, Supplementary Figure 4A-B**), confirming the utility of R-ATeam in co-imaging experiments with a kinase biosensor. Upon plotting the t_1/2_ of the mito-R-ATeam response against the t_1/2_ of the mito-ExRai-AMPKAR response to 2-DG or CCCP (**Supplementary Figure 4C**), we identified distinct response characteristics for both treatments. The mito-ExRai-AMPKAR response to CCCP treatment was consistent and rapid, regardless of differences in t_1/2_ of the mito-R-ATeam response to CCCP (**Supplementary Figure 4C**). Changes in ATP levels and AMPK activity following 2-DG treatment, however, were more gradual than the switch-like behavior observed with CCCP treatment (**Supplementary Figure 4C**, slope of best fit line from simple linear regression 2-DG, 0.14, CCCP, 2.01). There seems to be a graded AMPKAR response to 2-DG treatment, with longer t_1/2_ of R-ATeam correlating to longer t_1/2_ of AMPKAR. This may indicate a more sensitive coupling between AMPK energy sensing at the mitochondria and inhibition of oxidative phosphorylation compared to glycolysis. Follow up studies of cytosolic and nuclear AMPK activity in response to inhibition of glycolysis or oxidative phosphorylation may illuminate the distinct effects of these two energy production pathways on subcellular AMPK activity. Ultimately, further co-imaging applications of these two sensors can reveal more about the dynamic interplay between ATP levels and AMPK at subcellular resolution.

**Figure 4.**
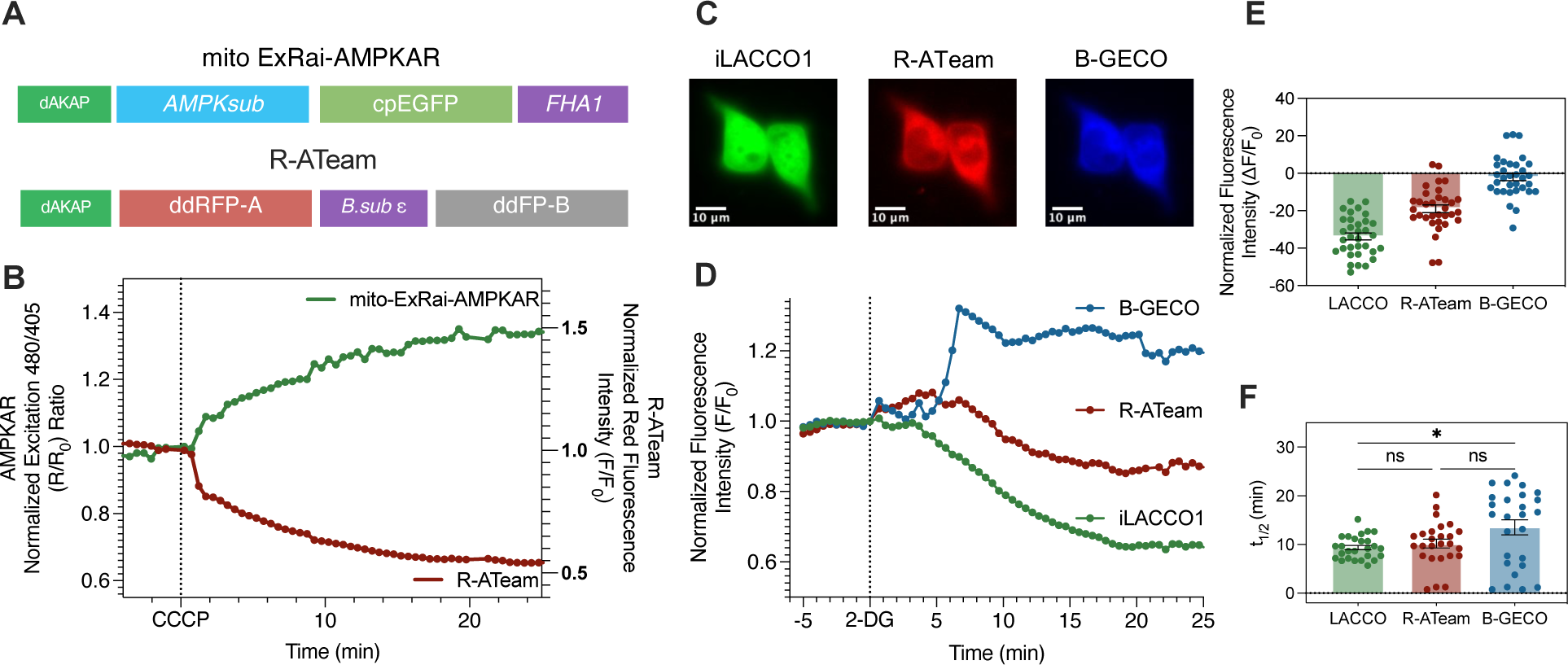
Multiplexed imaging of ATP with AMPK activity reporter (AMPKAR) and other cellular analytes. **(A)** Domain structures of mitochondria targeted mito-R-ATeam (top) and mito-ExRai-AMPKAR (bottom). **(B)** Representative single-cell time course traces of normalized red fluorescence intensity of R-ATeam (red, right axis) and normalized excitation ratio (480/405) of mito ExRai-AMPKAR (green, left axis) after CCCP treatment in HeLa cells co-expressing the two biosensors. Data is representative of three independent experiments. **(C)** Fluorescence images of representative HEK293T cells expressing iLACCO1 (left image), R-ATeam (middle image), and B-GECO (right image). Scale bar = 20 µm. **(D)** Representative single-cell time course traces of normalized intensity changes of B-GECO (blue, BFP), R-ATeam (red, RFP), and iLACCO1 (green, GFP) after 2-DG treatment. **(E)** Summary of 2-DG (n = 33) induced iLACCO1 (green), R-ATeam (red), and B-GECO (blue) responses from three independent experiments. **(F)** Summary of 2-DG (n = 33) induced iLACCO1 (green), R-ATeam (red), and B-GECO (blue) t_1/2_ from three independent experiments. One-way ANOVA with Dunnett’s multiple comparisons was used. ns (not significant), p > 0.05; *, p < 0.05.

Following these interesting observations on the complex interplay between ATP levels and AMPK activity, we aimed to achieve 3-color multiplexed imaging of key cellular analytes that are metabolically interconnected: ATP, lactate, and calcium (Ca^2+^). Lactate is a byproduct of glucose metabolism and lactate fermentation is used to produce ATP in the absence of oxygen^35^. On the other hand, a low cellular Ca^2+^ concentration can promote ATP metabolism^36^. Reduction of ATP levels by 2-DG has been reported to increase cellular Ca^2+^ levels^37^. However, the complex interplay between these analytes is still relatively under-studied. By co-expressing three single-color biosensors, R-ATeam, iLACCO1, and B-GECO^38^, we aimed to study the dynamics of ATP, lactate, and calcium induced by 2-DG treatment in HEK293T cells (n = 28) (**Figure 4C-F**). Upon 2-DG treatment, we consistently observed a decrease in green fluorescence of iLACCO1, correlating to a decrease in lactate, and a decrease in red fluorescence of R-ATeam, showing a decrease in ATP levels. We identified three distinct populations of cells with differing B-GECO responses to 2-DG (**Figure 4D, Supplementary Figure 4G-J**). One population of cells exhibited a sustained increase in calcium (n = 7) over a period of approximately 15-20 minutes (**Figure 4D, Supplementary Figure 4I**). Another population of cells did not show significant increases in Ca^2+^, as probed by B-GECO (n = 7), following 2-DG treatment (**Supplementary Figure 4G**). The third set of cells exhibited a significant (∼20-25%) but transient increase in B-GECO response (n = 19) following 2-DG treatment (**Supplementary Figure 4H**). B-GECO response (**Figure 4E**) and t_1/2_ (**Figure 4F**) was more variable compared to iLACCO1 and R-ATeam even when considering only the responding cells (sustained or transient responses). To further understand whether these three populations of cells also exhibited differences in ATP and lactate depletion, we conducted grouped analyses of t_1/2_ and response amplitude for each sensor. Overall, biosensor response amplitudes and t_1/2_ were not clearly grouped by transient or sustained B-GECO responses (**Supplementary Figure 4K**). The transient B-GECO response to 2-DG was faster than the iLACCO1 and R-ATeam responses in the same cells, but not significantly different in cells with sustained B-GECO response (**Supplementary Figure 4M**). Interestingly, iLACCO1 exhibited a faster response to 2-DG in the population of cells exhibiting no B-GECO response compared to cells with transient B-GECO response (**Supplementary Figure 4L**, none, 7.1 ± 0.9 min, transient. 9.0 ± 0.5 mins, p-value < 0.05). Such divergent calcium responses can be correlated to distinct lactate depletion characteristics because of simultaneous imaging of multiple sensors in single cells. Through these findings, we demonstrate that R-ATeam is suitable for multiplexed imaging experiments and characterizing cell heterogeneity in response to 2-DG.

## Discussion

Genetically encoded fluorescent protein-based biosensors of ATP have been essential to illuminating the real-time dynamics of ATP regulation in live cells. Biosensors such as ATeam1.03 and iATPsnFR have been used to elucidate cell-type specific metabolic differences and interactions^39–42^, the spatiotemporal correlation between ATP and other signaling molecules like calcium, sodium, and insulin^43–46^, and previously undefined roles of ATP in processes such as viral replication and cytopathic effect^47, 48^. These sensors have also been used to image changes in extracellular ATP concentrations, enabling characterization of ATP as a signal mediating cell-to-cell communication^13, 49, 50^. ATP sensors exhibiting various affinities to ATP have even been used to measure ATP concentrations in cortical astrocytes^51^. These previous studies highlight the importance of having a broadly applicable toolbox of sensors to gain novel insights on this key cellular analyte. In this study, we present an expanded palette of ATP sensors that exhibit improvements in dynamic range, signal to noise ratio, and response time. The utility of these sensors has been demonstrated in this study through characterization of subcellular ATP dynamics and multiplexed imaging of ATP with other cellular analytes and signaling activities.

We first presented an improved yellow/cyan FRET-based ATP biosensor, eATeam, which yielded a 40% improved response compared to ATeam1.03 in HEK293T cells. eATeam also had an improved signal to noise ratio. Another differentiator of eATeam from ATeam1.03 is the significantly faster response when monitoring 2-DG induced ATP decreases, which implies faster unbinding of ATP from the sensor and may be a more accurate indicator of real-time ATP dynamics. These distinctions make eATeam a valuable addition to the ATP reporting toolbox.

Subcellular ATP dynamics have not been thoroughly characterized. Many studies utilizing fluorescent biosensors of ATP focus on monitoring mitochondrial ATP levels^8, 9, 43^. Work has also been done to understand changes in ATP levels at the cell edge during cytoskeletal restructuring^52^ and nuclear ATP production in breast cancer cells^10^. In the original study introducing ATeam1.03, subcellularly targeted ATeam variants were used to define differences in basal yellow/cyan ratios of the sensor in the cytosol, mitochondria, and nucleus. Here, we take that work a step further using subcellularly targeted eATeam to visualize real-time changes in ATP levels in the cytosol, nucleus, mitochondria, and plasma membrane. The significance of ATP spatial regulation is highlighted at the beginning of its lifecycle when it is produced. Whereas glycolysis occurs in the cytoplasm near mitochondria, oxidative phosphorylation takes place across the inner mitochondrial membrane. ATP generated by oxidative phosphorylation in the mitochondrial matrix is transported out to the cytosol by mitochondrial ADP/ATP carriers^53^. Consistent function of these mitochondrial carriers is required to keep cytosolic ATP levels stably high. Although this may seem like a limiting factor for the delivery of subcellular ATP, we found that inhibiting oxidative phosphorylation in fact led to faster ATP depletion compared to inhibiting glycolysis in all probed compartments (**Figure 2E, J, O, T**), possibly due to the larger impact that oxidative phosphorylation has on cellular ATP production. A recent study identified distinct changes in mitochondrial and cytosolic ATP under hypoxia by co-expression of mitochondrial and cytosolic MaLionR and MaLionG, respectively, using a construct they named smacATPi^54^. Taking this previous study and our findings into consideration, it would be interesting to probe ATP dynamics in the mitochondrial matrix using an inner mitochondrial membrane targeted ATeam to interrogate the kinetics of ATP generation by oxidative phosphorylation and subsequent export of ATP from mitochondria. We revealed distinct spatiotemporal dynamics of ATP depletion upon inhibition of glycolysis or oxidative phosphorylation. This raises interesting questions about mechanisms of ATP delivery to cellular compartments from two different energy producing processes.

It is still unclear how cytosolic ATP, whether generated by glycolysis or oxidative phosphorylation, is transported to other subcellular compartments. In cardiac cells, nuclear ATP has been reported to accumulate by adenylate kinase-catalyzed phototransfer in a manner specifically coupled to oxidative phosphorylation, but not glycolysis^26^. Although we currently do not understand exactly how ATP produced by glycolysis is supplied to the nucleus, we observed that nuclear ATP was dependent on both glycolysis and oxidative phosphorylation (**Figure 2P-T**). In addition, nuclear ATP was depleted at a similar rate to cytosolic ATP upon inhibition of oxidative phosphorylation, but at a slower rate upon inhibition of glycolysis (**Supplementary Figure 2B**). This finding, building upon previous reports, suggests distinct mechanisms of ATP delivery to the nucleus depending on the process of production and presents another interesting direction for future study. Ultimately, we prove the utility of the improved FRET-based eATeam for probing subcellular ATP dynamics in live cells and present several interesting directions for future studies of subcellular ATP.

Expanding the toolbox of single-color ATP sensors can facilitate multiplexed imaging to unravel the network of ATP-dependent dynamic cellular signaling activities. The previously developed MaLion series introduced red, green, and blue ATP sensors incorporating circularly permutated fluorescent proteins as the reporting unit. We add to this existing toolbox of ATP sensors two spectrally distinct single-color ATP sensors incorporating dimerization-dependent FPs: R-ATeam (ddRFP-based) and G-ATeam (ddGFP-based). The response of R-ATeam and MaLionR induced by 2-DG treatment in HEK293T cells was comparable (**Supplementary Figure 3D**). One distinction is that the ddRFP (from R-ATeam) fluorescence spectrum is slightly further red-shifted (λ_ex_/λ_em_ = 585/610)^27^ compared to mApple (from MaLionR) (λ_ex_/λ_em_ = 565/585)^12^. This may be an important consideration in the design of multiplexed imaging experiments to reduce spectral overlap. R-ATeam performed similarly to eATeam with a 38.5 ± 14.0 % improved response compared to ATeam1.03. G-ATeam, on the other hand, exhibited a nearly 150 ± 14.3 % increase in response to 2-DG relative to ATeam1.03. Using these newly developed sensors, we imaged changes in nuclear and mitochondrial ATP levels following perturbation of glycolysis (2-DG) or oxidative phosphorylation (CCCP). By visualizing nuclear and mitochondrial ATP levels in the same cell simultaneously, we were able to uncover distinct subcellular ATP dynamics. We observed that mitochondrial ATP depletion was significantly faster than nuclear ATP depletion upon inhibition of glycolysis. However, mitochondrial and nuclear ATP were depleted at a similar rate when oxidative phosphorylation was disrupted (**Figure 3I, M**). Combined with the slower nuclear ATP depletion we observed upon glycolytic inhibition (via 2-DG) using Nuc-eATeam (**Figure 2T**), this data hints at differential relevance of these energy producing pathways to nuclear ATP supply. Such co-imaging studies can be applied to imaging inner and outer mitochondrial ATP changes, as mentioned above, to elucidate ATP generation and mitochondrial ATP export dynamics. Another interesting direction would be to co-image a plasma membrane localized ATP biosensor with an extracellular ATP biosensor to characterize cellular ATP export and import. Further studies utilizing single-color ATP sensors like R-ATeam and G-ATeam can reveal more about the spatiotemporal regulation of ATP.

The physiological relevance of studies on subcellular ATP dynamics can be enriched by concurrent imaging of ATP with other cellular analytes and signaling activities that are closely interwoven. Due to the direct relevance of ATP levels to AMPK activity, we co-imaged mitochondrial ATP levels (R-ATeam) and mitochondrial AMPK activity (AMPKAR). There were interesting differences between the dynamics of AMPKAR and R-ATeam responses to 2-DG and CCCP. CCCP induced consistently rapid activation of AMPK and depletion of ATP, resulting in a more switch-like effect of turning AMPK on or off. 2-DG, on the other hand, induced a more gradual and variable response from both biosensors. One potential explanation for this involves drug action. In addition to sensing energy status, AMPK is a redox sensor of mitochondrial reactive oxygen species (ROS)^55^. Since CCCP inhibits oxidative phosphorylation by uncoupling mitochondrial electron transport, it can lead to accumulation of ROS that, in turn, can help activate AMPK more rapidly than depletion of ATP alone. Harkening back to our results on variable calcium levels in response to 2-DG treatment (**Supplementary Figure 4G-I**), there may also be a connection between calcium response characteristics and AMPK activation kinetics upon 2-DG treatment. AMPK is regulated by two major upstream kinases, liver kinase B1 (LKB1) and calcium/calmodulin-dependent protein kinase kinase 2 (CAMKK2). Our previous study reported that mitochondrial and lysosomal AMPK activity were CAMKK2-dependent^7^. Regulation of AMPK by CAMKK2 is calcium-dependent^56^, so it stands to reason that mitochondrial AMPK may be sensitive to the variations in calcium induced by 2-DG treatment. Ultimately, we captured differences in response dynamics of AMPKAR and R-ATeam simultaneously within the same cell, and gleaned insight into the relationship between AMPK activity and ATP changes induced by CCCP and 2-DG. Future studies imaging subcellular AMPK activity, calcium, and ATP levels simultaneously in wild-type and CAMKK2 KO cells will be useful for characterizing the dynamic interplay between ATP, calcium, and AMPK.

We also successfully performed three-fold multiplexed imaging of calcium (B-GECO), ATP (R-ATeam), and lactate (iLACCO1), three cellular analytes that are metabolically related but do not have a well-established relationship. Although 2-DG induced intracellular calcium was previously reported^37^, it was not characterized temporally. We identified three distinct populations of cells defined by their B-GECO (Ca^2+^) response to 2-DG treatment: no response, transient response, and sustained response. By directly correlating B-GECO response characteristics to iLACCO1 and R-ATeam in the same cells, we further identified differences in lactate and ATP depletion in each subpopulation. In the population of cells exhibiting a transient calcium response, the B-GECO response precluded the R-ATeam and iLACCO1 responses. Interestingly, lactate depletion was slower in the cells with a sustained increase in calcium compared to cells without any detectable change in intracellular calcium (**Supplementary figure 4L**). A previous study linked extracellular lactate presence to inhibition of calcium transients in myocytes^57^, but a reverse relationship has not been characterized. Our findings may suggest a role for calcium in either promoting production or inhibiting depletion of lactate. Overall, these results demonstrate the advantage of multiplexed imaging experiments over single-parameter imaging for gaining insights on the interplay between signaling molecules to inform future investigations. The single-color ATP sensors presented here, R-ATeam and G-ATeam, are valuable tools that can be used to explore the complex interplay between multiple cellular processes.

In summary, we present an expanded palette of ATP sensors that can be utilized to gain insight into subcellular ATP dynamics and further investigate the convergence of ATP and other cellular signaling pathways.

## Materials and Methods

### Reagents and Constructs

2-deoxyglucose (2-DG, Sigma, D6134-250MG) was dissolved in 1x DPBS. Carbonyl cyanide m-chlorophenyl hydrazone (CCCP, Fisher Scientific, AC228131000) was dissolved in DMSO. FuGENE HD (E2311) was purchased from Promega. Mito-ExRai-AMPKAR (#192448), ATeam1.03 (#51958), MaLionRed (#113908), B-GECO1 (#32448) are all available on addgene. iLACCO1 (intracellular lactate sensor) was kindly provided by the Campbell lab^58^.

To optimize ATeam1.03, we tested 3 different cyan/yellow fluorescent protein pairs present in other constructs: mTurquoise2 and cpmVenus[E172] (from pRSETb-AKAR)^20^, mCerulean and cpVenus[E172] (from pRSETb-AKAR4)^19^, and mCerulean and YPet (from pRSETb-CANAR2)^18^. We generated the ddRFP-based R-ATeam from pRSETb-Rab-ICUE^59^. Primers were designed to generate fragments of the different pairs of fluorescent proteins and ATP sensing subunit of ATeam1.03 with overlapping regions to facilitate fragment joining by Gibson assembly. Through this strategy, we generated ATP sensor variants incorporating different fluorescent protein pairs in pRSETb backbones. These were then subcloned into the pcDNA3 vector by digestion with SacI and PaeI (SphI) (ThermoFisher FD1134 and ER0601) and ligated using T4 DNA ligase from New England Biolabs (M0202S). The ddGFP-based G-ATeam was cloned using pcDNA3-dPlcR as the template^60^. We designed primers to incorporate the ddGFP and FP-B fragments from pcDNA3-dPIcR and overlap with the ATP sensing subunit of ATeam1.03. For all Gibson assemblies, the ampicillin resistance gene was split into two fragments to serve as an internal control for successful assembly. Primers used for generating these constructs are listed below:

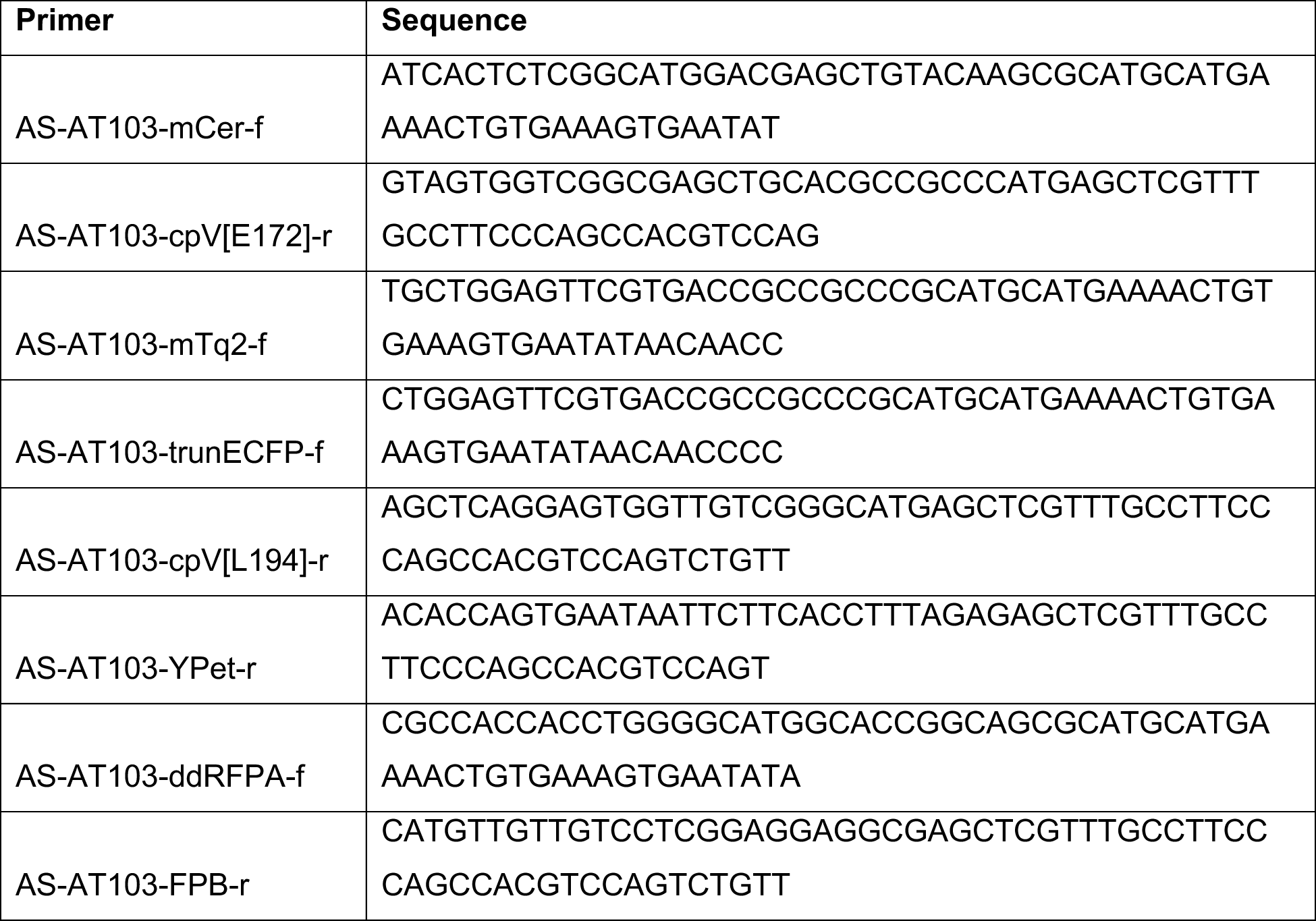
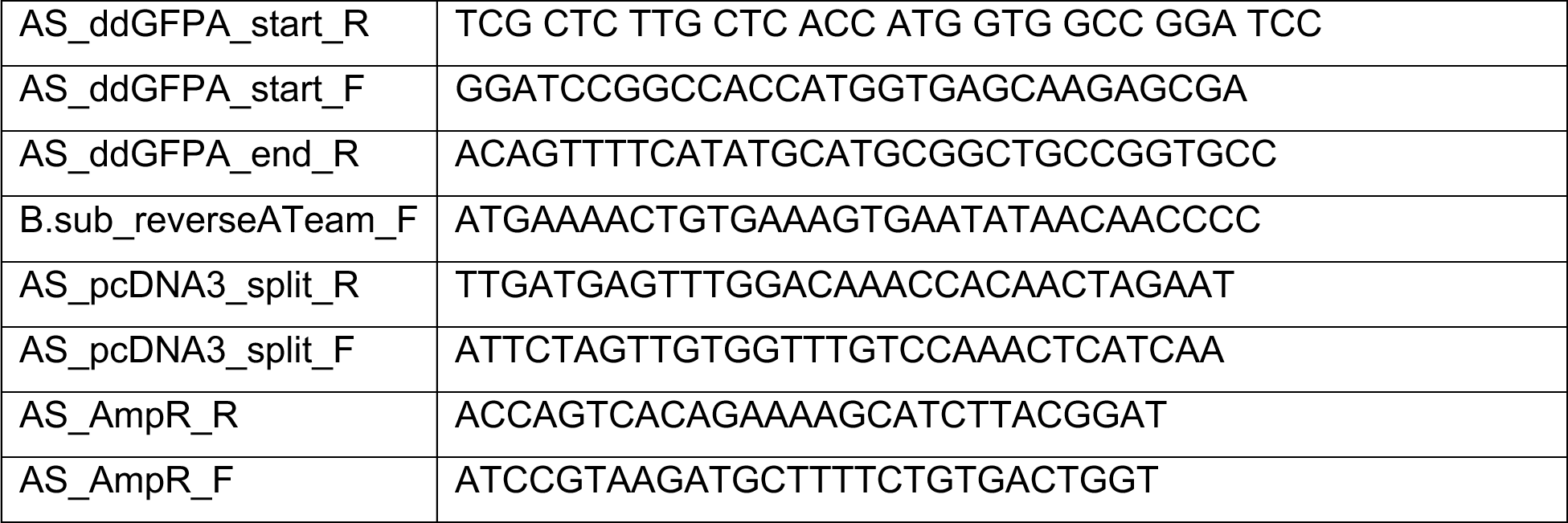

Subcellularly targeted ATP biosensors were generated by subcloning BamHI/XbaI (Thermofisher FD1464 and FD0275, respectively) digested biosensors into vector backbones containing the correct localization sequences (on the N-terminus), listed below:

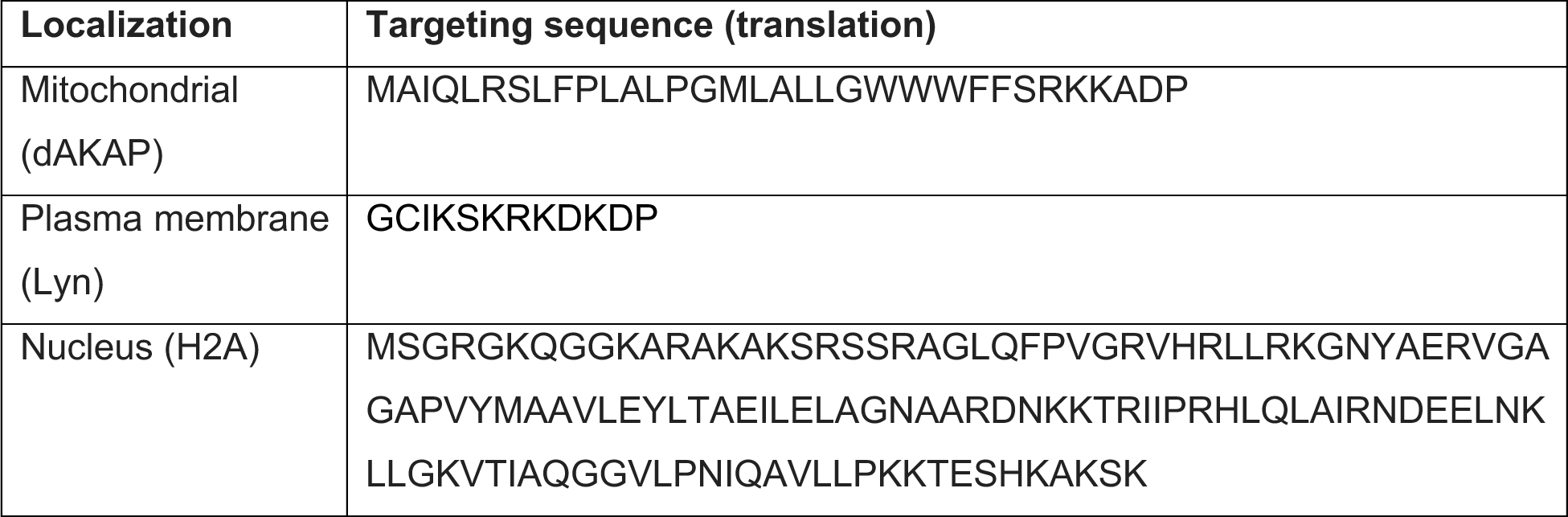

All generated constructs were verified by sequencing.

### Cell culture and transfection

HEK293T, MEF and HeLa cells used in this study were acquired from ATCC. HEK293T and MEF cells were cultured in Dulbecco’s modified Eagle medium (DMEM; Gibco 11995-065) with 4.5 g/L glucose, 10% fetal bovine serum (FBS, Gibco 26140-079), and 1% (v/v) penicillin/streptomycin (Pen/Strep, Gibco 15140-122). HeLa cells were cultured in Dulbecco’s modified Eagle medium (DMEM; Gibco 11885-084) with 1g/L glucose, 10% fetal bovine serum (FBS, Gibco 26140-079), and 1% (v/v) penicillin/streptomycin (Pen/Strep, Gibco 15140-122). For live-cell imaging experiments, HeLa cells were plated onto 35-mm glass-bottomed dishes (CellVis D35-14-1.5-N) and transfected in serum-free DMEM 2-24 hours after plating. HEK293T cells were seeded onto 35-mm glass-bottom dishes coated with Poly-D-Lysine (Gibco, A3890401), and transfected in serum-free DMEM 2-24 hours after plating. All cells were transfected using FuGENE HD (Promega, E2311) and imaged 16-24 hours after transfection.

### Fluorescence imaging

Before imaging, cells were washed once with imaging medium (1X Hank’s balanced salt solution with 2g/L glucose, pH 7.4, made from 10X HBSS (Gibco, 14065)) and incubated in imaging medium for 10-30 minutes at 37°C before imaging. Imaging was conducted in the dark at room temperature. Dishes were imaged for 30 minutes, once every 30 seconds. At least 5 images were obtained before treatment to establish the baseline response. Treatments of 2-DG (40 mM) or CCCP (5 nM) were added at the indicated times. A Zeiss Axio Observer Z7 microscope (Carl Zeiss) equipped with a 40x/1.4NA oil objective, a Definite Focus 2 system (Carl Zeiss), and a photometricsPrime95B sCMOS camera (Photometrics) was used to acquire images. The microscope was controlled by the MATScope imaging suite (GitHub, https://github.com/jinzhanglab-ucsd/MatScopeSuite), which utilizes open-source software: MATLAB (Mathworks) and µmanager (Micro-Manager). For Ateam1.03 and eATeam imaging, dual emission ratio imaging was performed using a 420DF20 excitation filter, 455DRLP dichroic mirror, a 475DF24 emission filter for CFP, and a 535DF25 emission filter for YFP. For R-ATeam, RFP was imaged using a 572DF35 excitation filter, a 594DRLP dichroic mirror, and a 645DF75 emission filter. To image G-ATeam or iLACCO1, GFP was imaged using a 480DF20 excitation filter, a 505DRLP dichroic mirror, and a 535DF50 emission filter. To image mito-ExRai-AMPKAR, two excitation filters (480DF20 and 405DF20), a 505DLRP dichroic mirror, a 535DF50 emission filter was used. To image B-GECO, a 405DF20 excitation filter, a 455DLRP dichroic mirror, and a 475DF24 emission filter was used. An LEP MAC6000 control module (Ludl Electronic Products Ltd) was used to alternate filter sets. Exposure times were 50-500ms, with images being taken of relevant channels once every 30 seconds.

### Image analysis and calculations

Images obtained by the MATScope imaging suite were analyzed by a custom MATLAB code available on Github. For untargeted constructs, whole cells were randomly selected for analysis. For targeted (subcellular) imaging, regions of interest (ROI) were selected in the cytosol, around mitochondria, at the plasma membrane, or in the nucleus. Background fluorescence was used to correct raw fluorescence intensities. For FRET-based eATeam and ATeam1.03, the yellow/cyan FRET emission ratios were calculated after background-subtraction of each channel. For R-ATeam, the background-subtracted red emission intensity was obtained. For G-ATeam and iLACCO1, the background-subtracted green emission intensity was obtained. For mito-ExRai-AMPKAR, the cpGFP excitation ratio was calculated using background-subtracted GFP emission after 480nm or 405 nm excitation (480/405). For B-GECO, the blue emission intensity was obtained. All emission ratios or intensities were normalized to the event-point, which was the time right before drug addition (R_0_ or F_0_). Normalized emission ratios or intensities were used to calculate response, t_1/2,_ and signal to noise ratio. Response (ΔR/R_0_ or ΔF/F_0_) was calculated by subtracting the average of the baseline (timepoints before drug addition) from the average of the last 10 timepoints and then dividing it by the average of the baseline: (R_mean(t50-60)_ – R_mean(t1-10)_)/ R_mean(t1-10)_. Change in Normalized Emission Ratio (ΔR/R_0_) reported in **Figure 1** and **Supplementary Figure 1** for ATeam1.03 and eATeam was calculated using baseline corrected values. The time to half response (t_1/2_) of each cell was calculated by identifying the time at which half the endpoint response was reached. The signal to noise ratio (SNR) was calculated by dividing the response of each cell by the standard deviation of the baseline (before addition of drug).

### Statistics and reproducibility

Data was compiled in Microsoft Excel, and figures and statistical analyses were prepared using GraphPad Prism 9. Unpaired student’s t-test (two-tailed) was used for comparing two parametric data sets, and paired student’s t-test (two-tailed) was used for comparing two-parametric data sets that were acquired simultaneously in the same cell. Ordinary one-way ANOVA with Dunnett’s multiple comparisons test was used to compare three or more data sets. P-values < 0.05 were defined as statistically significant. Ns (not significant) indicates a p-value > 0.05, * indicates a p-value between 0.01 and 0.05, ** indicates a p-value between 0.001 and 0.01, *** indicates a p-value between 0.0001 and 0.001, and **** indicates a p-value < 0.0001. P-value definitions were reported in the figure legend where relevant. The number of independent experiments that data is acquired from and number of analyzed cells (n) are reported in figure legends. Time courses are plotted with the mean response ± standard error of the mean, and dot plots are depicting mean ± SEM.

---

**Supplementary Figure 1.**
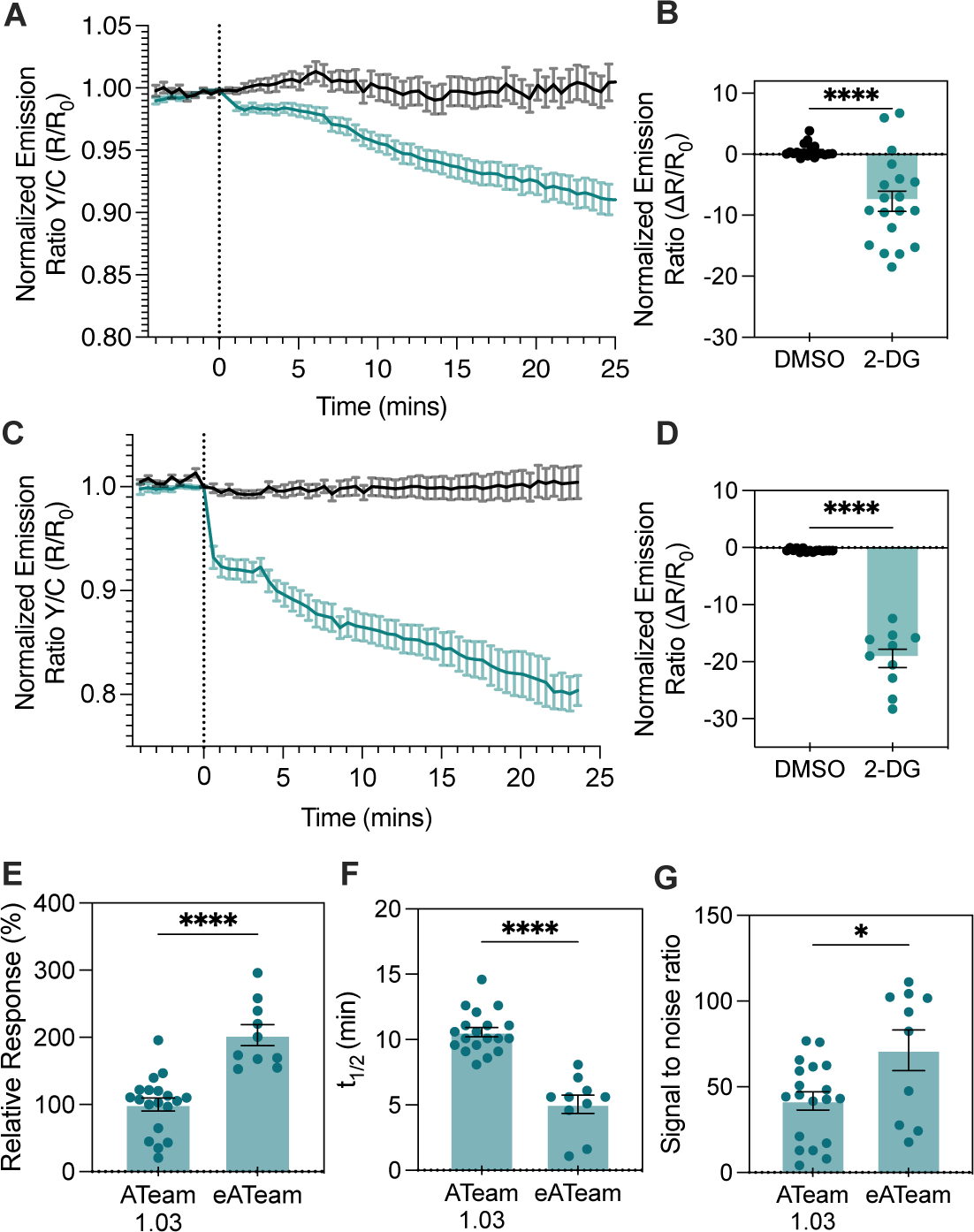
Comparison of ATeam1.03 and eATeam in MEF cells. **(A)** Averaged time course traces of normalized emission ratio (yellow/cyan) of ATeam1.03 in MEF cells treated with DMSO (black, vehicle control) or 2-DG (blue). Data were from at least three independent experiments. Shaded area indicates standard error of the mean (SEM). **(B)** Bar graph shows the average response 25 minutes after DMSO (n = 19) or 2-DG (n = 19) addition. Unpaired student’s t-test (two-tailed) is used. ***, p<0.001. **(C)** Averaged time course traces of normalized emission ratio (yellow/cyan) of eATeam in MEF cells treated with DMSO (black, vehicle control) or 2-DG (blue). Data were from least three independent experiments. Shaded area indicates standard error of the mean (SEM). **(D)** Bar graph shows the average response 25 minutes after DMSO (n = 16) or 2-DG (n = 10) addition. Unpaired student’s t-test (two-tailed) is used. ****, p < 0.0001. **(E)** Comparison of 2-DG induced responses of ATeam1.03 (n = 19) and eATeam (n=10) in MEF cells. Bar graph shows the average response time 25 minutes after 2-DG addition and error bars represent SEM. Unpaired student’s t-test (two-tailed) is used. ****, p < 0.0001. **(F)** Comparison of t_1/2_ of 2-DG induced responses of ATeam1.03 (n = 19) and eATeam (n=10). Bar graph shows the average t_1/2_ and error bars represent SEM. Unpaired student’s t-test (two-tailed) is used. ****, p < 0.0001.

**Supplementary Figure 2.**
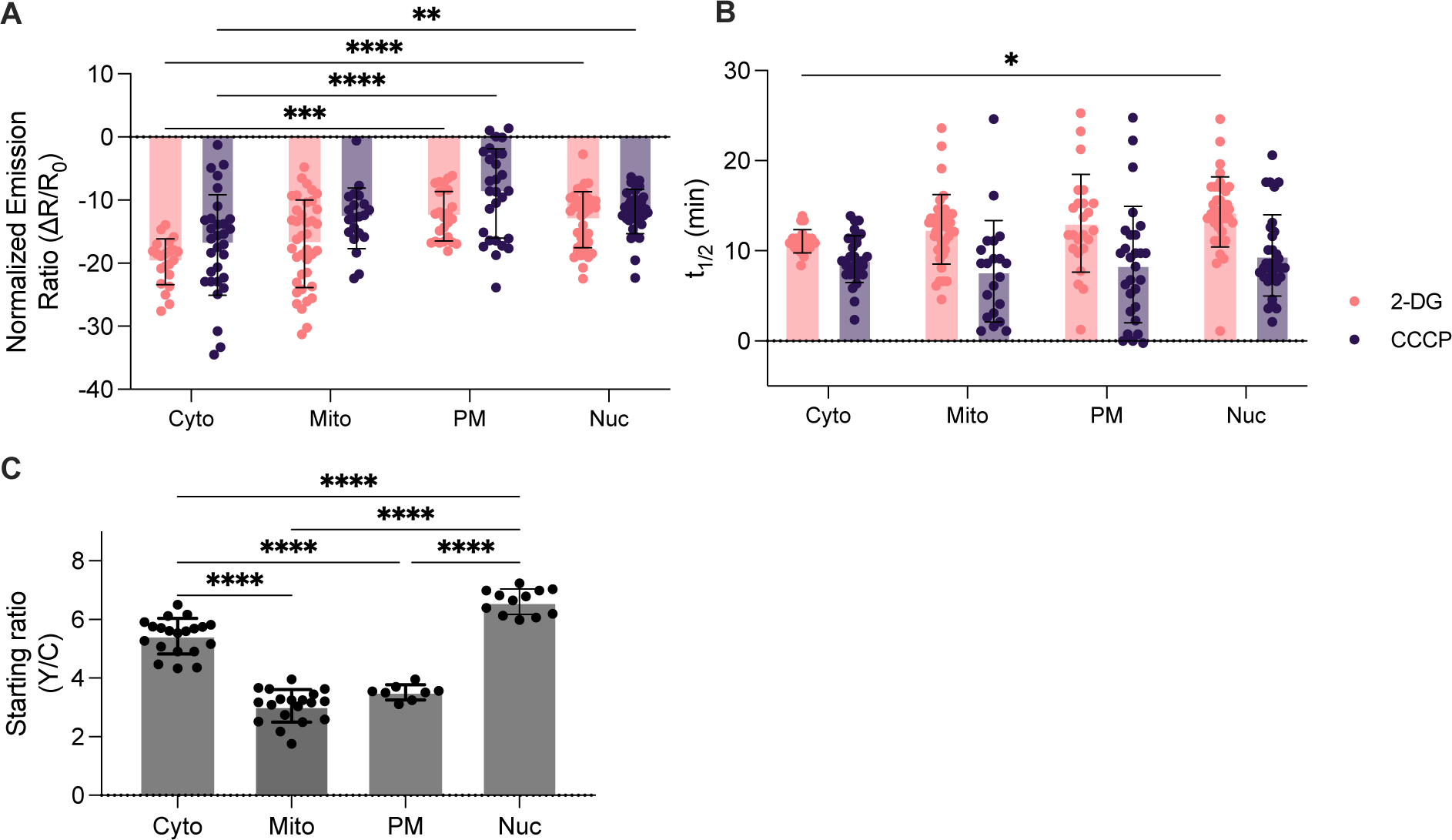
Comparison of subcellular eATeam responses to 2-DG. Summary of 2-DG (pink) and CCCP (purple) induced response **(A)**, t_1/2_ **(B)**, and starting ratios **(C)** of subcellular eATeams (cytosol, mitochondria, plasma membrane, and nucleus targeted) expressed in HEK293T cells. Data is from three independent experiments. One-way ANOVA with Dunnett’s multiple comparisons is used. ****, p < 0.0001, ***, p < 0.001, **, p < 0.01, *, p < 0.05.

**Supplementary Figure 3.**
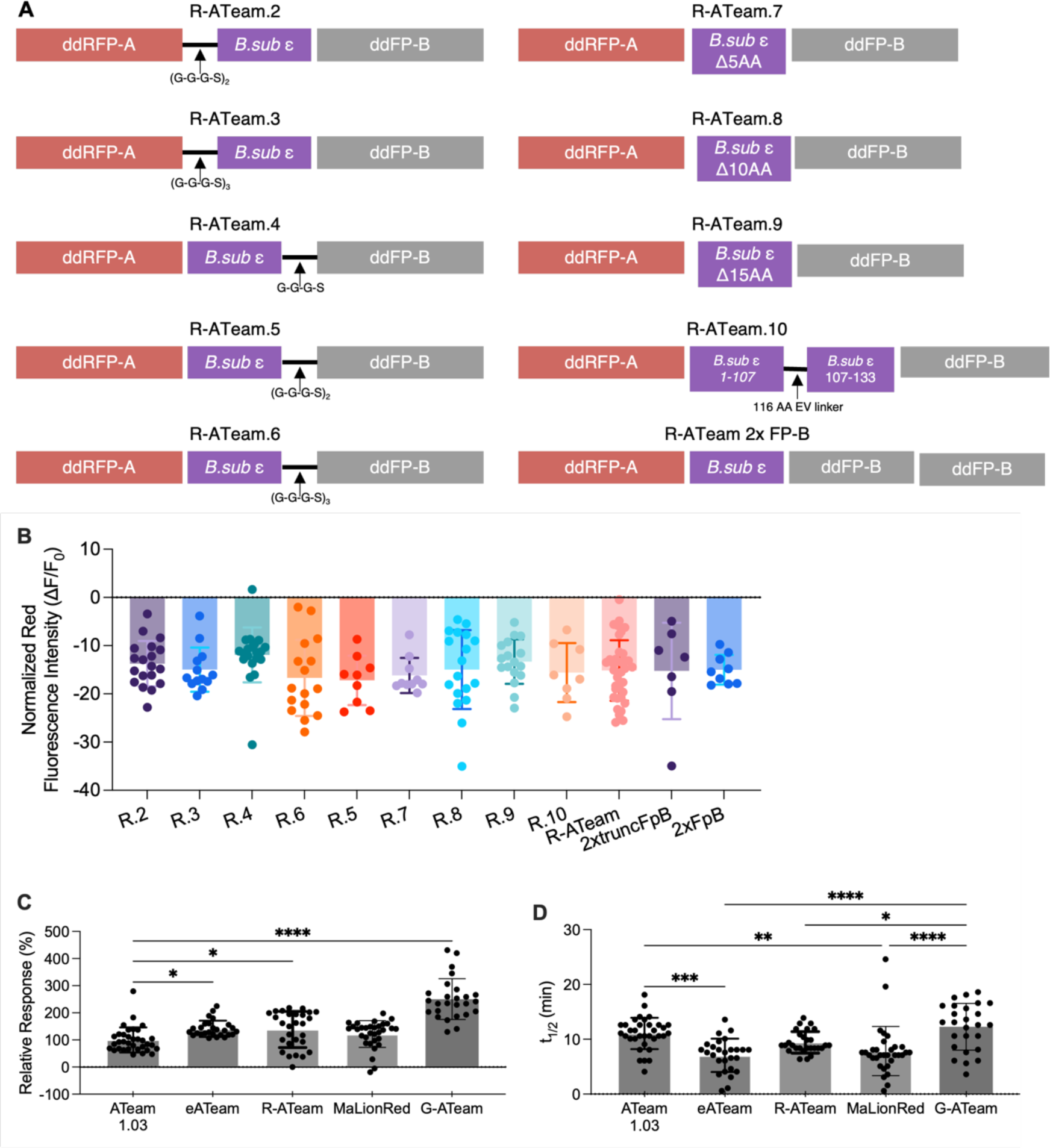
Characterization of different R-ATeam variants. **(A)** Domain structures of R-ATeam variants tested. **(B)** Summary of R-ATeam variants’ endpoint response induced by 2-DG. Bar graph shows the average response at time = 25 minutes, error bars indicate standard deviation. Oordinary one-way ANOVA with Dunnett’s multiple comparisons is used, no significance between any two variants’ responses. **(C)** Relative ATeam1.03 (n = 34), eATeam (n = 27), R-ATeam (n = 28), and G-ATeam (n = 26) response to 2-DG. Calculated by dividing each response value by the mean response of Ateam 1.03. Bar graph shows the average response at time = 25 minutes and error bars represent standard deviation. Ordinary one-way ANOVA with Dunnett’s multiple comparisons is used. ****, p < 0.0001, *, p< 0.05. **(D)** ATeam1.03 (n = 34), eATeam (n = 27), R-ATeam (n = 28), MaLionR (n = 32), and G-ATeam (n = 26) T one-half to 2-DG. Bar graph shows the average T one-half and error bars represent standard deviation. Ordinary one-way ANOVA with Dunnett’s multiple comparisons is used. ****, p < 0.0001, *, p< 0.05.

**Supplementary Figure 4.**
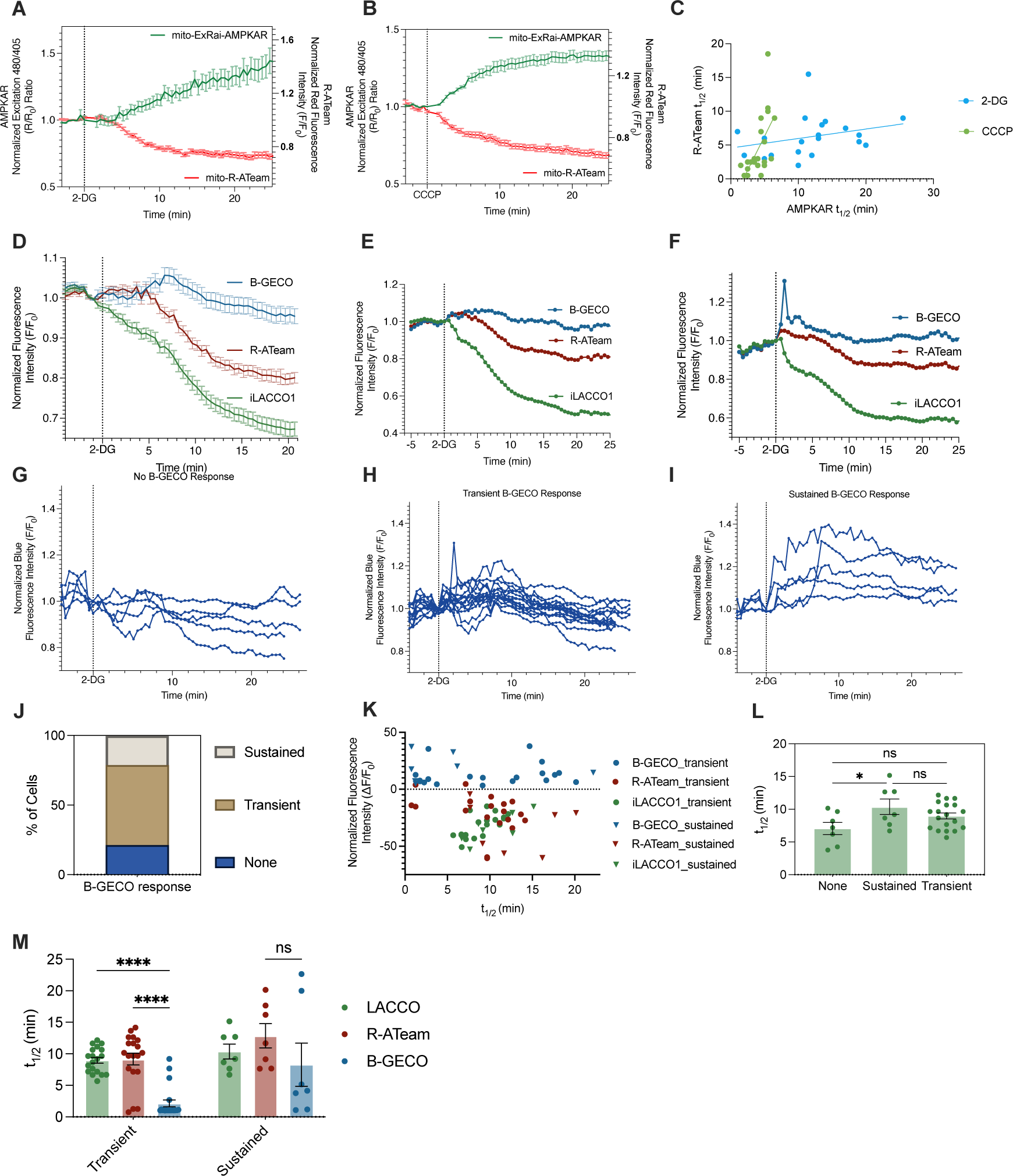
Average curves and more representative cell data for multiplexed imaging experiments. Averaged time course traces of normalized excitation ratio (480/405) of mito-ExRai-AMPKAR (green, left axis) and normalized red fluorescence intensity of R-ATeam (red, right axis) in HeLa cells treated with 2-DG (**A**) or CCCP (**B**). Data were from at least three independent experiments. Shaded area indicates standard error of the mean (SEM). **(C)** t_1/2_ of mito-R-ATeam response plotted against t_1/2_ of mito-ExRai-AMPKAR response after 2-DG (blue) or CCCP (green) treatment in the same cell. Best-fit lines following simple linear regression are plotted for each group (2-DG or CCCP). **(D)** Averaged time course traces of normalized fluorescence intensity of B-GECO (blue), R-ATeam (red), and iLACCO1 (green) induced by 2-DG treatment in HEK293T cells (n = 33). Data were from at least three independent experiments. Shaded area indicates standard error of the mean (SEM). (**E-F**) Representative single cell time course traces of B-GECO (blue), R-ATeam (red), and iLACCO1 (green) induced by 2-DG treatment in HEK293T cells. Cells represent variable responses to 2-DG by B-GECO found in the imaged population of cells. (**G-I**) Single cell time course traces of normalized blue fluorescence intensity of B-GECO in HEK293T cells after 2-DG treatment, grouped by response characteristic: (**G**) no response (n = 7), (**H**) transient response (n = 19), (**I**) and sustained response (n = 7). (**J**) Percentage of imaged cells that exhibited no (none, n = 7), sustained (n = 19), or transient (n = 7) B-GECO response to 2-DG. (**K**) Response plotted against t_1/2_ of iLACCO1, R-ATeam, and B-GECO response after 2-DG treatment by B-GECO response characteristic (transient or sustained response to 2-DG). (**L**) Comparison of t_1/2_ of iLACCO1 response to 2-DG in cells exhibiting no (none), sustained, or transient B-GECO response to 2-DG. Bar graph represents mean and error bars represent SEM.*, p < 0.05; ns, p > 0.05, indicates not significant. (N) Comparison of t_1/2_ of iLACCO1 (green), R-ATeam (red), or B-GECO (blue) response to 2-DG in cells exhibiting no (none), sustained, or transient B-GECO response to 2-DG. Bar graph represents mean and error bars represent SEM. Ordinary one-way ANOVA with Dunnett’s multiple comparisons test was used. ****, p < 0.0001.

